# DREAM: A Toolbox to Decode Rhythms of the Brain System

**DOI:** 10.1101/2020.01.29.926204

**Authors:** Zhu-Qing Gong, Peng Gao, Chao Jiang, Xiu-Xia Xing, Hao-Ming Dong, Tonya White, F. Xavier Castellanos, Hai-Fang Li, Xi-Nian Zuo

## Abstract

Rhythms of the brain are generated by neural oscillations across multiple frequencies, which can be observed with multiple modalities. Following the natural log linear law of frequency distribution, these oscillations can be decomposed into distinct frequency intervals associated with specific physiological processes. This perspective on neural oscillations has been increasingly applied to study human brain function and related behaviors. In practice, relevant signals are commonly measured as a discrete time series, and thus the sampling period and number of samples determine the number and ranges of decodable frequency intervals. However, these limits have been often ignored by researchers who instead decode measured oscillations into multiple frequency intervals using a fixed sample period and numbers of samples. One reason for such misuse is the lack of an easy-to-use toolbox to implement automatic decomposition of frequency intervals. We report on a toolbox with a graphical user interface for achieving local and remote decoding rhythms of the brain system (DREAM) which is accessible to the public via GitHub. We provide worked examples of DREAM used to investigate frequency-specific performance of both neural (spontaneous brain activity) and neurobehavioral (in-scanner head motion) oscillations. DREAM analyzed the head motion oscillations and found that younger children moved their heads more than older children across all five frequency intervals whereas boys moved more than girls in the age interval from 7 to 9 years. It is interesting that the higher frequency bands contains more head movements, and showed stronger age-motion associations but the weaker sex-motion interactions. Using the fast functional magnetic resonance imaging data from the Human Connectome Project, DREAM mapped the amplitude of these neural oscillations into multiple frequency bands and evaluated their test-retest reliability. A novel result indicated that the higher frequency bands exhibited more reliable amplitude measurements, implying more inter-individual variability of the amplitudes for the higher frequency bands. In summary, these findings demonstrated the applicability of DREAM for frequency-specific human brain mapping as well as the assessments on their measurement reliability and validity.

## 1 Introduction

Rhythms of the brain are generated by neural oscillations occurring across multiple frequencies [5]. The natural logarithm linear law (N3L) offers a theoretical framework for parcellating these brain oscillations into multiple frequency intervals linking to distinct physiological roles [22]. Remarkably, when graphed on the natural logarithm scale, the centers of each frequency interval fall on adjacent integer points. Thus, distances between adjacent center points are isometric on the natural logarithm scale, resulting in a full parcellation of the whole frequency domain where each parcel of the frequencies is fixed in theory, namely frequency intervals. These frequency intervals have been repeatedly observed experimentally [6]. This characteristic suggests that distinct physiological mechanisms may contribute to distinct intervals. Functional magnetic resonance imaging (fMRI), a non-invasive and safe technique with an acceptable trade-off between spatial and temporal resolution, has the potential to contribute to the study of certain neural oscillations in the human brain in vivo. In early fMRI studies of the human brain, researchers tended to treat oscillations across different frequencies without differentiation. Low-frequency oscillations measured by resting-state fMRI (rfMRI) have been assessed primarily in the frequency range of 0.01 to 0.1 Hz, a range in which spontaneous brain activity has high signal amplitude [4,20]. While such efforts have been somewhat informative, treating this broad frequency range in a unitary manner may conceal information carried by different frequency intervals. To address this issue, an early study decomposed the rfMRI signals into multiple frequency intervals using the N3L theory (Slow-5: 0.01 - 0.027 Hz, Slow-4: 0.027 - 0.073 Hz, Slow-3: 0.073 - 0.198 Hz, Slow-2: 0.198 - 0.25 Hz) [47]. This exploration demonstrated the feasibility of mapping distributional characteristics of oscillations” amplitude in both space and time across multiple frequency intervals in the brain.

Since then, an increasing number of rfMRI studies have employed such methods by directly applying these frequency intervals, and have detected frequency-dependent differences in brain oscillations in patients. Specifically, these differences were mostly evident between Slow-4 and Slow-5 amplitudes [14,16,19,21,28,42]. Such frequency-dependent phenomena have also been explored using other rfMRI metrics including regional homogeneity detected in the Slow-3 and Slow-5 frequency ranges [34]. While the lower and upper bounds of the frequency intervals are fixed in theory, their highest and lowest detectable frequencies and frequency resolution are determined by the sampling parameters (e.g., rate and duration) in computational practice. However, the above-mentioned studies applied the frequency intervals from earlier studies [8,47] rather than to use those matching their actual sampling settings. To address this situation, we developed an easy to use toolbox to decode the frequency intervals by applying the N3L theory. This toolbox, named DREAM, is based on MATLAB with a graphical user interface (GUI). Here, we introduce the N3L algorithm and its DREAM implementation. Neural oscillations reflected by the human brain spontaneous activity measured with resting-state functional MRI and head motion data during mock MRI scans were employed as two worked examples to demonstrate the use of DREAM to perform frequency analyses.

## 2 Methods and Algorithms

Neuronal brain signals are temporally continuous but they are almost always measured as discrete data for practical reasons. The characteristics of the sampled data should meet the criterion of the sampling theorem proposed by American electrical engineers Harry Nyquist and Claude Shannon. The core algorithm to determine the frequency boundaries of measured neuronal signals in DREAM is based on the Nyquist-Shannon sampling theorem. Specifically, per the theorem, sampling frequency and sampling time determine the highest and lowest frequencies that can be detected and reconstructed. Sampling data retains most of the information contained in the original signals if the sampling frequency is at least twice the maximum frequency of the continuous signals. As for neuronal signals, the highest frequency that could be detected and reconstructed is determined by the sampling frequency, or by the sampling interval which is equal to the reciprocal of the sampling frequency, as shown in formula (1)

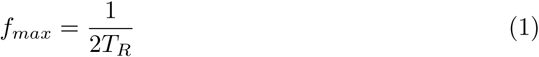

where *f*_*max*_ represents the highest frequency that could be detected in the neuronal signal and *T*_*R*_ represents the sampling interval.

The lowest frequency in neuronal signals that could be detected depends on the sampling time. As shown in formula (2), in order to distinguish the lowest frequency in neuronal signals, the sampling time should be equal to or larger than the reciprocal of two times the lowest frequency

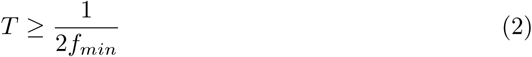

where *T* represents the sampling time, and *f*_*min*_ represents the lowest frequency in neuronal signals that could be distinguished.

Since the sampling time is equal to the number of samples multiplied by the sampling interval, the lowest frequency can be calculated by formula (3):

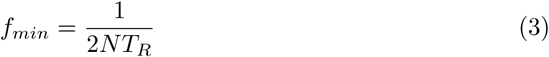

where *N* represents the number of samples.

According to the N3L theory, neural oscillations in mammalian brain formed a linear hierarchical organization of multiple frequency bands when regressed on a natural logarithmic scale. The center of each band would fall on each integer of the natural logarithmic scale (Fig. 1-1). Thus, adjacent bands have constant intervals that equals to one, which correspond to the approximately constant ratios of adjacent bands on the linear scale (Fig. 1-2). With the highest and lowest frequencies reconstructed, N3L can derive the number of decoded frequencies and the boundaries of each frequency interval (Fig. 1-3). Accordingly, when graphed on the natural log scale, the center of each decoded frequency is an integer. Thus, adjacent center points on the natural log scale are equidistant, which corresponds to the same proportion of adjacent center points” values on the linear scale. Based upon this theorem, after performing a linear regression analysis for the highest and lowest frequencies acquired previously, we can determine the central frequencies, as well as the number frequency intervals that can be decoded.

**Figure 1.**
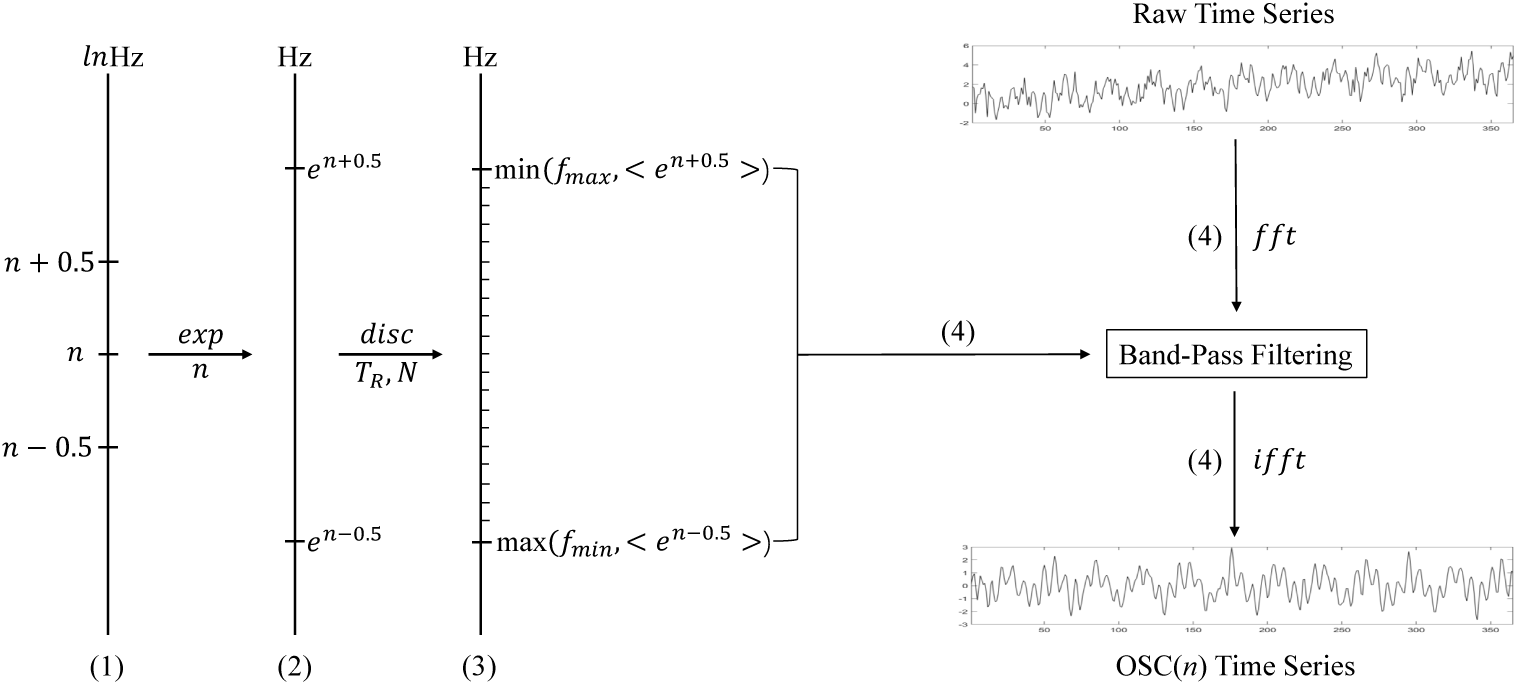
The flowchart on the DREAM algorithm. **(1)** N3L theory defines an oscillator with a length-one frequency band centered at *n*, i.e., OSC(*n*), in the natural log space. **(2)** In original frequency space, it expands the frequency band (**e**^*n*−0.5^, **e**^*n*+0.5^) Hz. **(3)** This frequency band can be discretized with a sampling procedure with *N* points and *T*_*R*_ rate in terms of the classical signal theory. **(4)** This computational frequency band is for a band-pass filtering process to extract the OSC(*n*) from the raw time series.

Finally, the decoding process integrated in DREAM performs band-pass filtering with the frequency intervals provided by DREAM in the previous steps (Fig. 1-4). This is implemented by the MATLAB built-in function **fft** and **ifft** to perform direct and inverse time-frequency transformation on the signals for individual decoded frequency intervals, respectively. All the above steps are illustrated as the flowchart in Figure 1.

### Interface and Usage

DREAM has been shared and released with the Connectome Computation System [36]. After downloading the package at GitHub, users will need to add the directory where the package is stored into the MATLAB path. The package can then be launched by entering “DREAM” in the MATLAB command line. DREAM integrates its GUI (two buttons) into its flash screen (Fig. 2).

**Figure 2.**
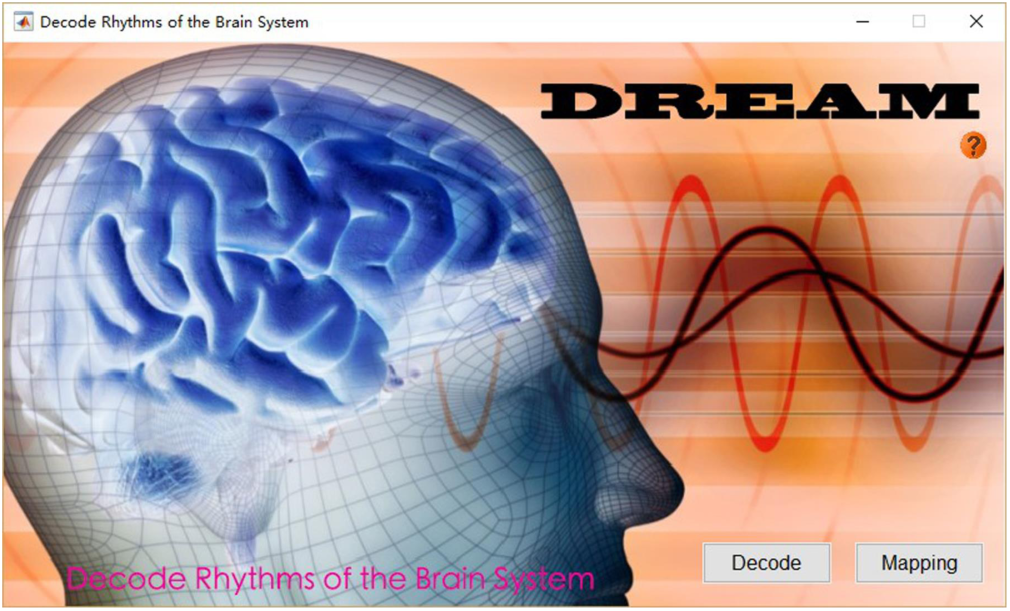
DREAM Flash.

### Program Interface

Users should enter or organize their data into the predefined directory structure (Fig. 3) before start processing the data. The **work directory** is where the subject directories are stored (full path). Individual data should be stored in each **subject directory** or a sub-folder inside (**data directory**). DREAM has a main interface (Fig. 4) for setting up the structure (the left side) and previewing the plots of time series from the data selected (the right side).

**Figure 3.**
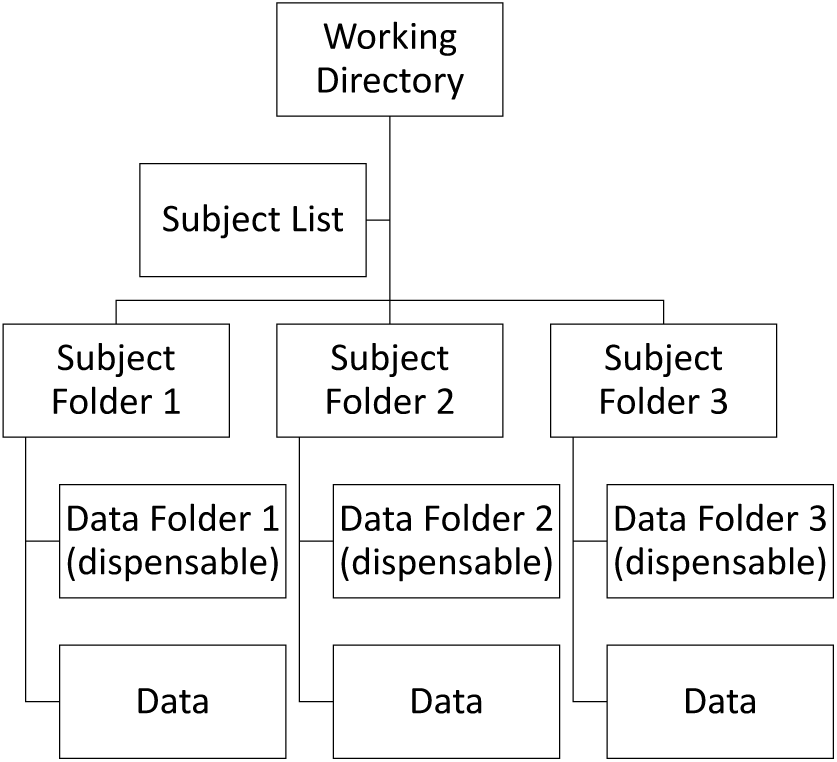
DREAM DirTree.

**Figure 4.**
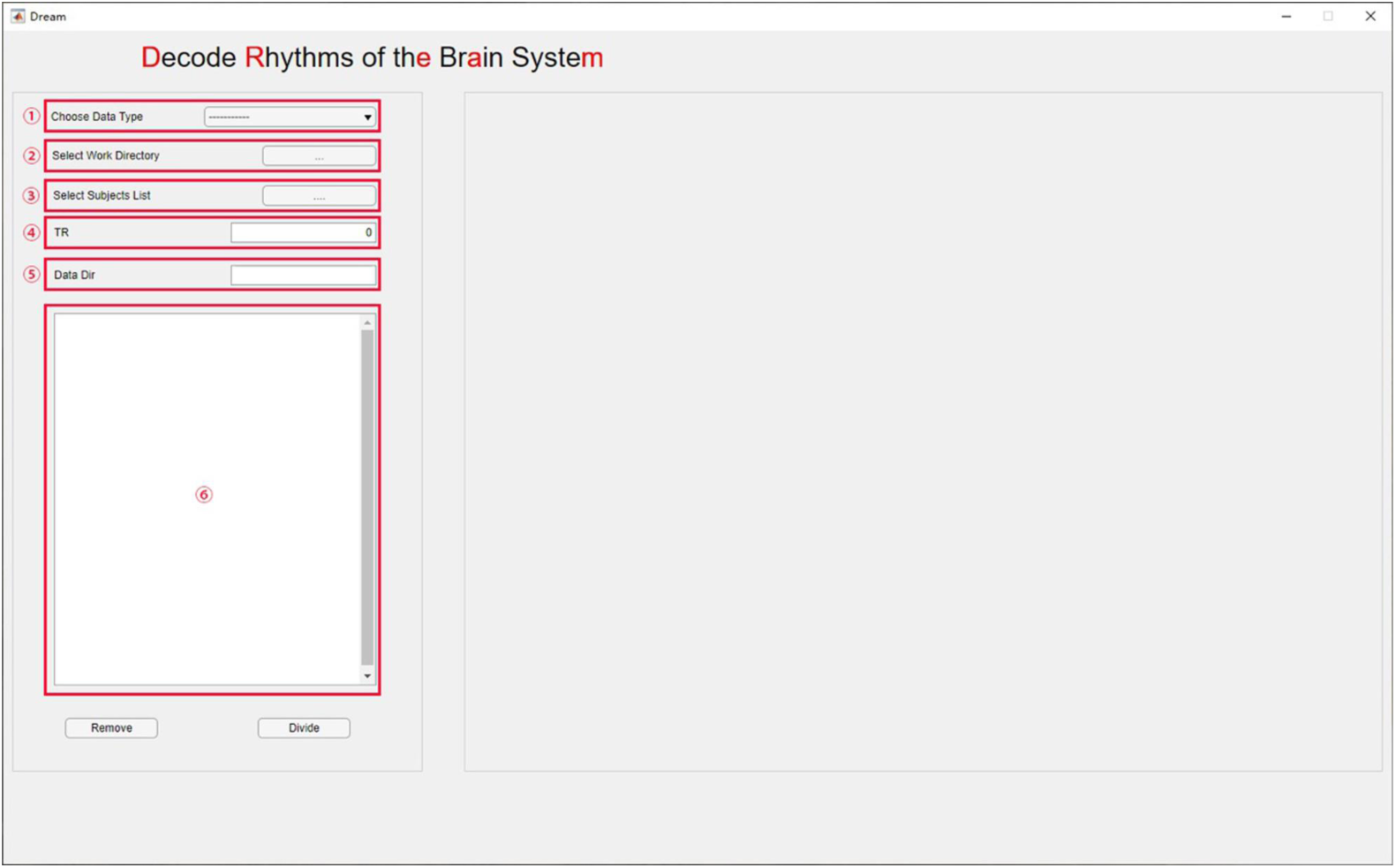
The main interface of DREAM.

### GUI Usage

We introduce how to use the graphical interface step by step in below. The circled numbers in Figure 4 correspond to the analyzing steps in this section.

- Step 1 - **select the data type**: Click the drop-down box to choose the data type to be analyzed.
- Step 2 - **set up the work directory**: Click the path selection button to set the work directory in the dialog box that pops up.
- Step 3 - **batch process**: Select the subject list file in the popped-up dialog box by clicking the file selection button.
- Step 4 - **set up sampling rate**: Enter the sampling interval in seconds (*T*_*R*_) in the input box (in some cases, this can be automatically extracted from the header information).
- Step 5 (optional) - **data directory**: If the data are stored in a sub-folder inside the subject directory, type the name of the data directory in the input box.

After all the above parameters are set up, data meeting the requirement will appear in the list-box (Fig. 4-6), from where the user can remove unwanted data by selecting the file name and clicking the **Remove** button. Finally, by clicking the **Divide** button, a user can start the decoding program. The outputs contain a set of decoded files and a **csv** file that records the boundary frequencies of each decoded band. The outcomes can be directly used for subsequent analyses.

## 3 DREAM1: Frequency-dependent oscillations of in-scanner head motion in 3-16 years-old children

In-scanner head motion has been treated as a confound in fMRI studies, especially in studies of children and patients with psychiatric disorders. Many studies have shown the effects of motion on fMRI results such as increases of short-distance correlations and decreases of long-distance correlations in rfMRI-derived connectivity metrics [23,27,38]. Researchers have proposed various methods to correct motion effects in fMRI studies. In contrast, studying head motion as a neurobehavioral trait has long been overlooked (see an exception in [41]), especially in children. Here, we use DREAM to quantify head motion data acquired from preschool and school children in a mock scanner using a novel multi-frequency perspective. We hypothesized that: 1) head motion is a behavioral trait associated with age; 2) there are sex differences in head motion in children; and 3) the head motion effects are frequency-dependent.

### Participants and Data Acquisition

We recruited 94 participants (47 females) between 3 to 16 years of age as part of the Chinese Color Nest Project [39,48], a long-term (2013-2022) large-scale effort on normative research for lifespan development of mind and brain (CLIMB) [9]. All participants were from groups visiting during the **Public Science Open Day of the Chinese Academy of Sciences**, with the approval of at least one legal guardian. The experiment was performed in a mock MRI scanner at the site of the MRI Research Center of the Institute of Psychology, Chinese Academy of Sciences. The mock scanner was built by PST (Psychology Software Tools, Inc.) using a 1:1 model of the GE MR750 3T MRI scanner in use at the institute. It is used for training young children to lie still in a scanner before participating the actual MRI scanning session. It is decorated with cartoon stickers to provide a children-friendly atmosphere. Head motion data were acquired with the MoTrack Head Motion Tracking System (PST-100722). The system consists of three components: a MoTrack console, a transmitter and a sensor. The sensor is worn on the participant”s head and provides the position of the head relative to the transmitter. For each participant, head motion is displayed on the computer screen in real-time. The original sampling rate of the system is 103 Hz. The averaging buffering size is 11 samples, which results in a recording sampling rate of 9.285 Hz. The participants were instructed to rest quietly on the bed of the mock scanner for around three and half minutes without moving their heads or bodies. They were watching a cartoon film inside the scanner during the “scanning” to simulate movie-watching scanning. The data acquisition period was designed to resemble the real MRI scanning environment, with a recording of scanning noises of the real MRI machine played as the background noise.

### Data Analysis

Head motion data are recorded in text files consisting of six parameters for each time point, three translation (millimeters) and three rotation (degrees) measures. The first three parameters are displacements in the superior, left and posterior directions, respectively. The last three parameters are rotation degrees in the three cardinal rotational directions. We converted the original data into frame-wise displacement (FD), a single parameter scalar quantity representing head motion proposed by Power and colleagues [23]. To correct for spikes caused by sudden movements, which may bias mean FD values, we applied the AFNI **3dDespike** command (version 17.3.06) to the FD time series. Data without this preprocessing was also analyzed and supported reproducible patterns. Then time-windows were determined and applied before feeding the data into DREAM. We retained 1672 sampling points from the zeroed time point (time point when the original six parameters were set to zero), which equaled a duration of three minutes. After preprocessing, we used DREAM to decode the data. Of note, the original FD values were all positive. After decoding, the time series of decoded bands were demeaned, which means the average values of all decoded time series were very near to zero. Thus, we took the absolute value of decoded frequency intervals to calculate mean FD values, which were used in subsequent statistical analyses. Inspired by many human growth curves modeled by exponential function and the scatter plots on the head motion data, we first converted the head motion data using the natural logarithm transformation and then assess the relationship between FD and age by using linear regression models to fit the FD data in each frequency interval with age. We conducted this regression for boys and girls, respectively, and tested whether the slopes and intercepts are significantly different between boys and girls. Of note, this method is equivalent to an Analysis of Covariance (ANCOVA) [29]. These analyses were also applied to the standard deviation of FD time series to test the stability of head motion.

### Results

Six participants were excluded from further data analysis due to sampling periods less than three minutes. Another four participants were excluded because their mean FD values were three standard deviations higher than the mean value of the whole group (i.e., outliers). Total 42 boys (age: 3 - 14 years, 8.7 ± 3.0) and 42 girls (age: 4 - 16 years, 8.4 ± 3.1) were included in our final analyses. No significant differences in age were found between males and females. All the findings derived with the head motion data without **despike** preprocessing are highly similar to those of using **despike**, which are reported as following. Meanwhile, all the results derived from the linear regression models are replicated by the ANCOVA model.

#### Frequency Decomposition

Since all the head motion data have the same sampling frequency and sampling period, DREAM decoded all the FD time series into the same six frequency intervals named according to [6] (Slow-4: 0.033 to 0.083 Hz, Slow-3: 0.083 to 0.22 Hz, Slow-2: 0.222 to 0.605 Hz, Slow-1: 0.605 to 1.650 Hz, Delta: 1.650 to 4.482 Hz, Theta: 4.482 to 4.643 Hz). This theta band is too narrow comparing with its full range (up to 10 Hz) to be reliable for the analyses, and thus not included in our analyses. The full band and the five frequency bands from an individual child are depicted in Figure 5 and Figure 6.

**Figure 5.**
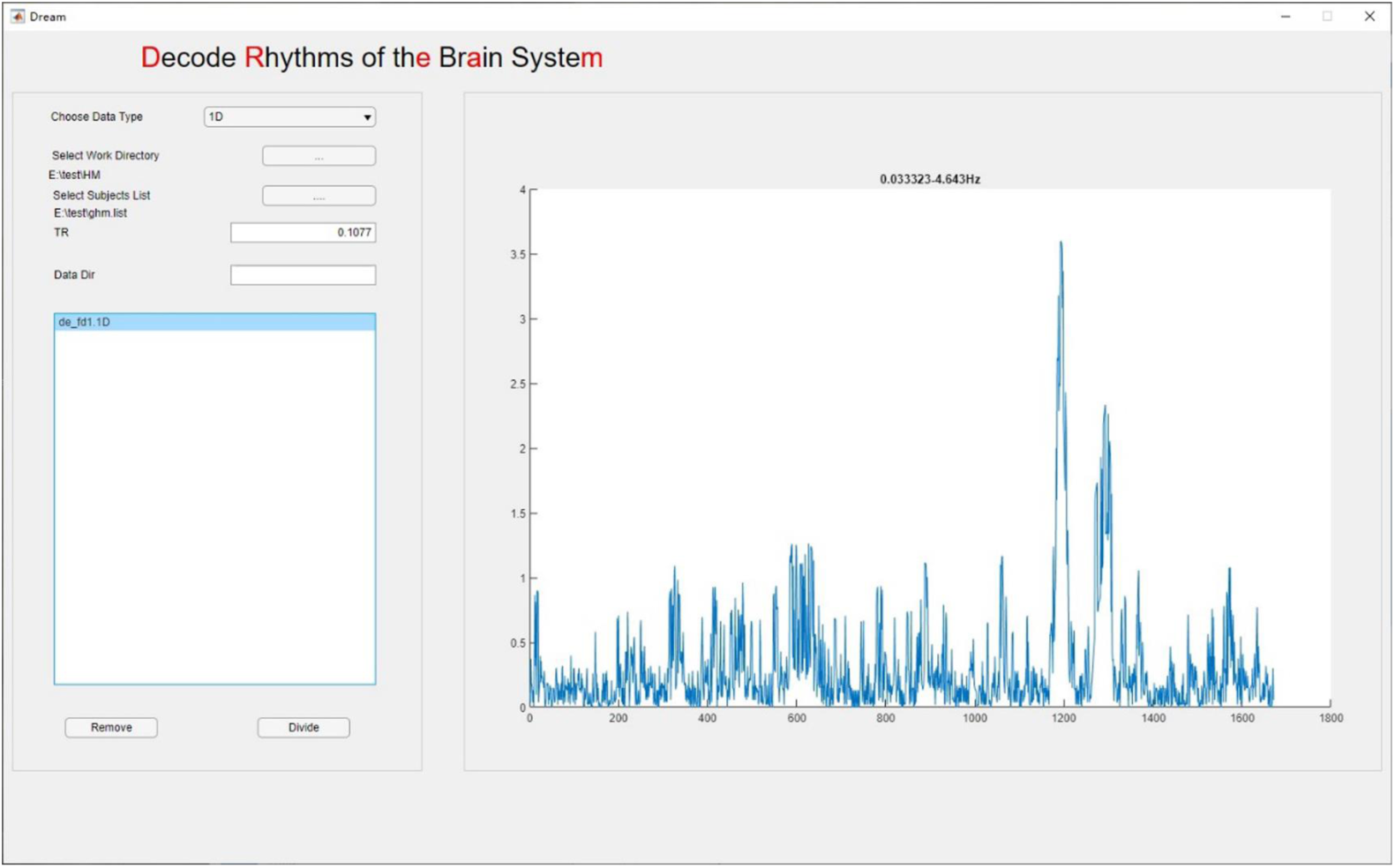
A preview of the original FD time series from a participant.

**Figure 6.**
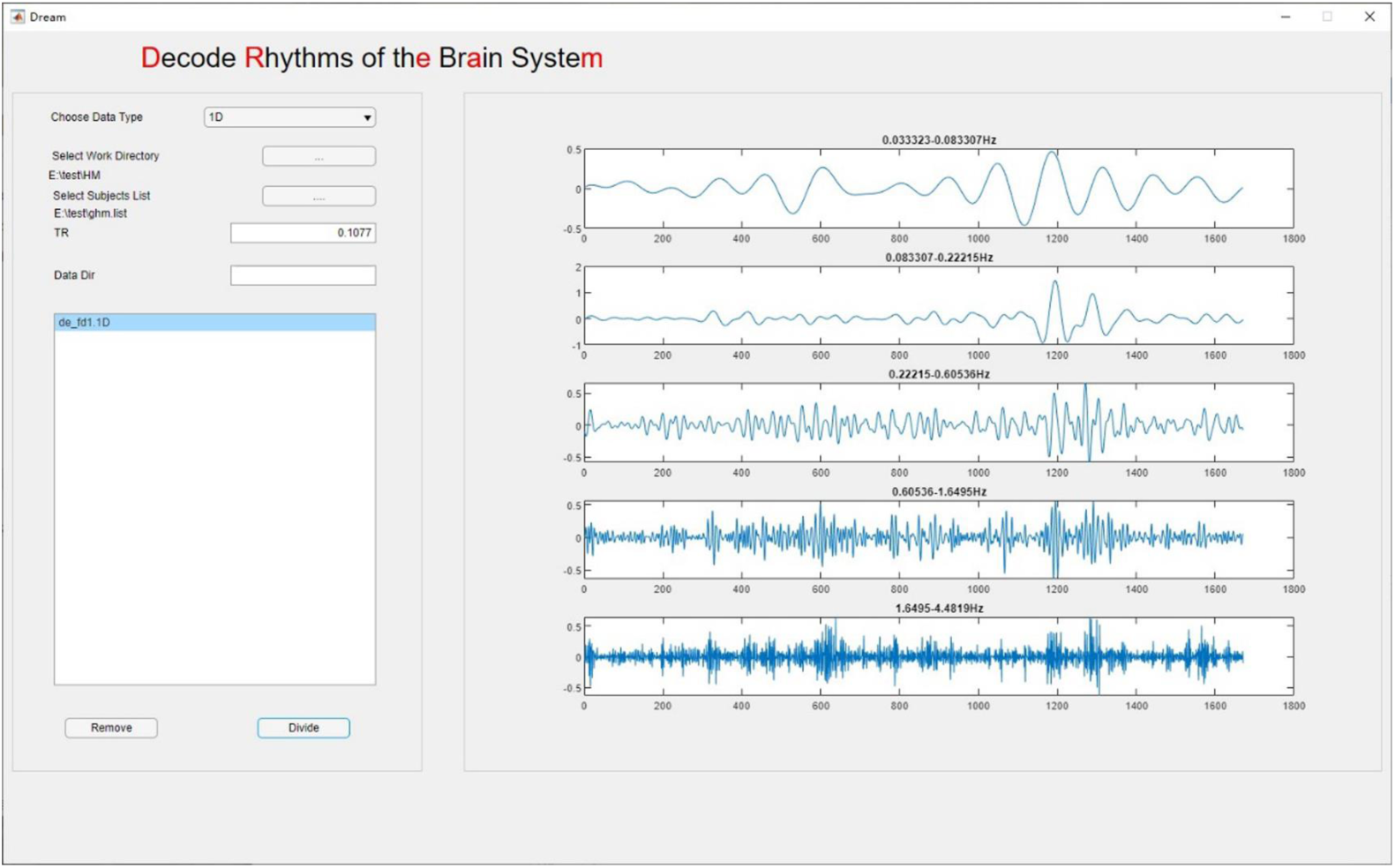
DREAM decodes FD time series into the five bands.

#### Age-related Head Motion Changes across Frequencies

Results from the linear regression analysis yielded significant negative correlations between age and mean FD values across all the five bands for both boys and girls (*df* = 40, FDR corrected *p* < 0.05):

- Slow-4: boys, *p* = 0.018, *R*^2^ = 0.218; girls, *p* = 0.034, *R*^2^ = 0.195
- Slow-3: boys, *p* = 0.008, *R*^2^ = 0.249; girls, *p* = 0.027, *R*^2^ = 0.203
- Slow-2: boys, *p* = 0.001, *R*^2^ = 0.314; girls, *p* = 0.017, *R*^2^ = 0.221
- Slow-1: boys, *p* < 0.001, *R*^2^ = 0.358; girls, *p* = 0.013, *R*^2^ = 0.230
- Delta : boys, p < 0.001, R2 = 0.380; girls, p = 0.008, R2 = 0.250

The relationship between age and mean FD values are plotted in Figure 7, indicating that younger children tend to move more than older ones, and this trait correlation held in both boys and girls. We also performed a similar linear regression analysis between the standard deviations of decoded FD values and age, and observed similar outcomes that the standard deviations were significantly negatively correlated with age across frequency bands and sexes. This showed older children are more stable with their head motion than younger children.

**Figure 7.**
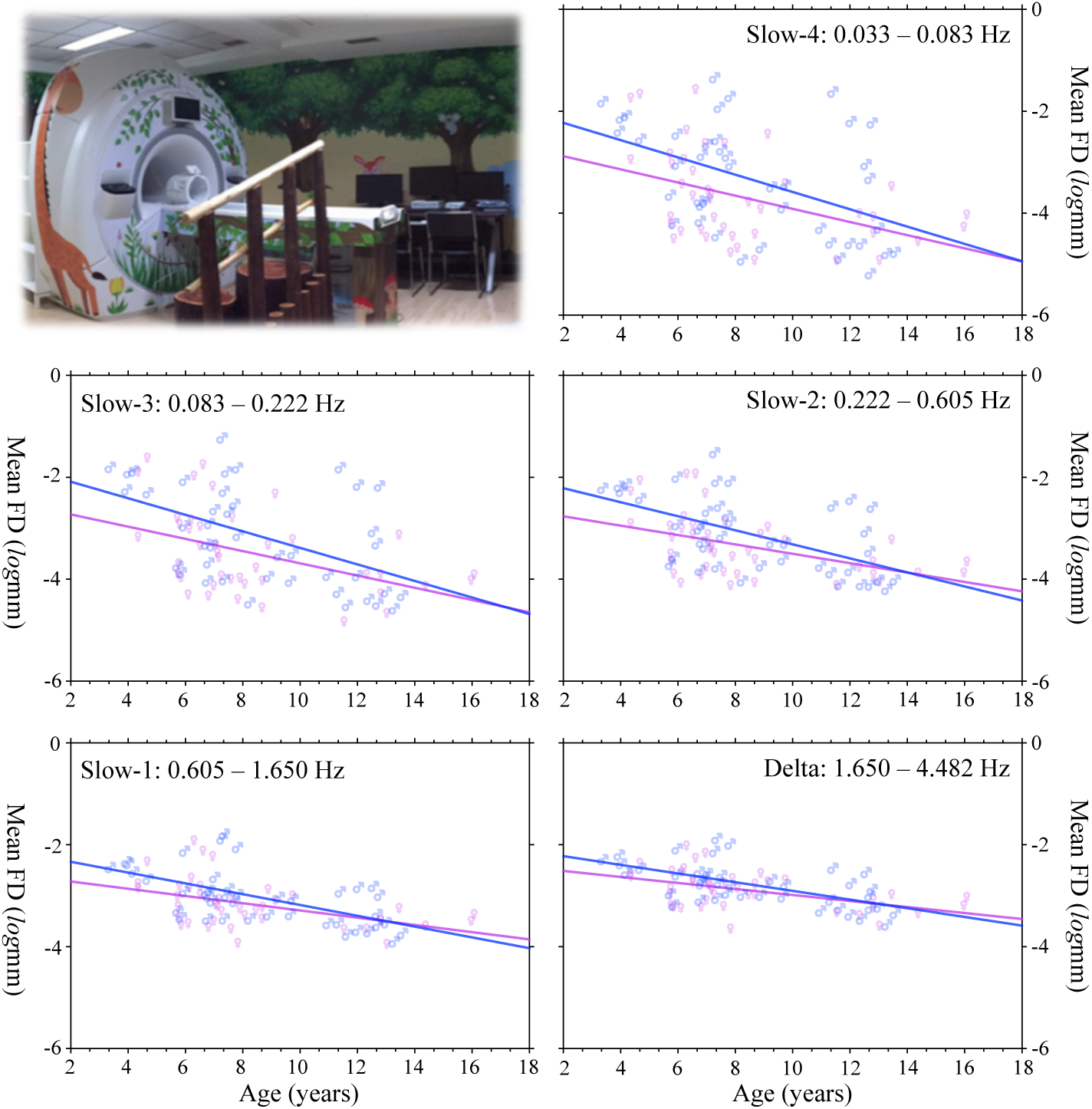
Nonlinear age-motion relationship across the five frequency bands. The plots are based upon the log transformed motion data, indicating the exponential growth model 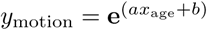. The upper-left panel shows the Mock scanning facility in the Magnetic Resonance Imaging Research Center at the Institute of Psychology, Chinese Academy of Sciences.

We further tested if the two lines are different between boys and girls. Statistical results revealed no such sex-related effect (*df* = 80, FDR corrected *p* > 0.05):

- Slow-4: slope, *p* > 0.5, *F* = 0.383; intercept, *p* = 0.494, *F* = 3.979
- Slow-3: slope, *p* > 0.5, *F* = 0.531; intercept, *p* = 0.385, *F* = 4.428
- Slow-2: slope, p > 0.5, F = 1.177; intercept, p = 0.486, F = 4.010
- Slow-1: slope, *p* > 0.5, *F* = 1.326; intercept, *p* = 0.849, *F* = 3.042
- Delta : slope, *p* > 0.5, *F* = 1.222; intercept, *p* = 0.968, *F* = 2.822

Inspired by the trend that sex-related differences in mean FD are smaller in higher frequency bands, especially evident for early stages, we thus divided all the participants into three age groups (3 to 6 years: 14 boys, 18 girls; 7 to 9 years: 14 boys, 15 girls; 10 to 16 years: 14 boys, 9 girls) and compared mean FD values between males and females in each age group using two-way (sex and frequency band) ANOVA with repeated measures. Figure 8 summarized the results of an increasing pattern of head motion from slow to fast bands for all the age groups (3-6yrs: *F* (4) = 10.90, *p* = 1.65 × 10^−7^; 7-9yrs: *F* (4) = 20.62, *p* = 1.20 × 10^−12^; 10-16yrs: *F* (4) = 23.95, *p* = 3.06 × 10^−13^). Meanwhile, we observed a significant interaction between sex and frequency band in 7 to 9 years old children (*F* (4, 1) = 3.22, *p* = 0.0154) but not for the other groups (3-6yrs: *F* (4, 1) = 0.195, *p* = 0.940; 10-16yrs: *F* (4, 1) = 1.065, *p* = 0.380).

**Figure 8.**
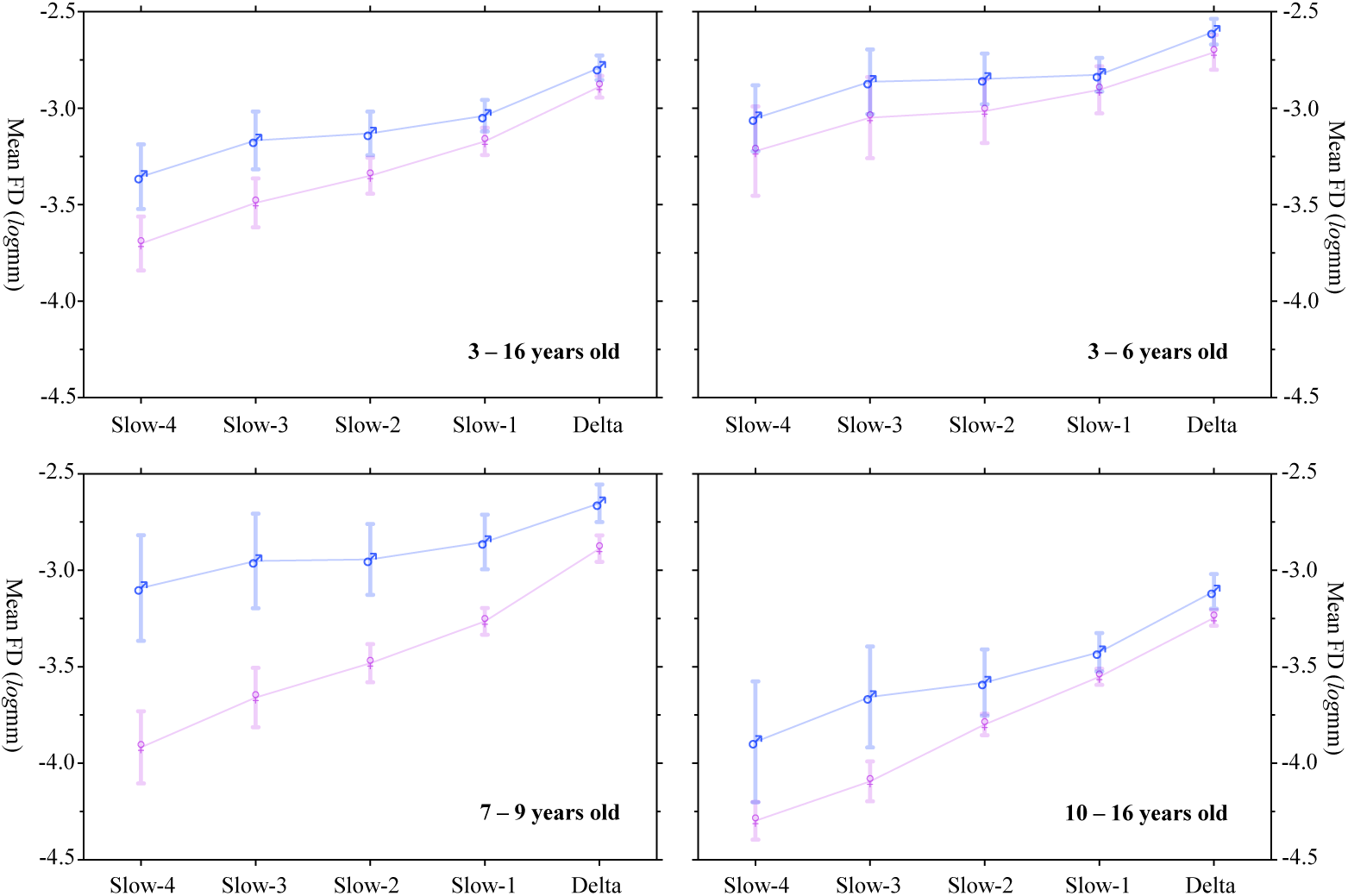
Sex-frequency interactions on head motion across ages. All the participants (3 to 16 years old) are divided into three age groups: 3 to 6 years, 7 to 9 years, 10 to 16 years). A two-way (sex and frequency band) ANOVA with repeated measures compares mean FD values between males and females in each age group.

## 4 DREAM2: Frequency-dependent spatial ranking and reliability of low-frequency oscillations

The amplitude of low frequency fluctuation (ALFF) is a common metric used in fMRI studies that reflects regional amplitude of the signal intensity”s fluctuations in a frequency range [40]. Previous studies revealed variations of ALFF in both spatial and frequency domains in the resting-state brain. From the perspective of spatial distribution, in the typical resting-state frequency range (e.g., 0.01-0.1 Hz), the neural oscillations showed higher ALFF in grey matter than white matter [4,32]. It is noted that ALFF reaches its peaks in visual areas [17], posterior structures along brain midline [4,44] and in cingulate and medial prefrontal cortices [12]. In frequency domain, BOLD oscillations distributed to grey matter were mainly in Slow-4 and Slow-5, while its white matter oscillations were dominated by Slow-3 and Slow-2 [47]. Specifically, higher ALFF in Slow-4 was detected in the bilateral thalamus and basal ganglia whereas the slow-5 oscillators exhibited higher ALFF in the ventromedial prefrontal cortex, precuneus and cuneus (replicated in [37]). These findings revealed the frequency-specific characteristics of resting-state ALFF. The previous studies are limited by their sampling precision (*T*_*R*_ ≤ 2000ms), and studies on the ALFF distribution across more accurate bands and their reliabilities are still lacking. For examples, the Slow-2 frequency band derived in [47] has quite small overlap with its theoretical range and thus may limit both reliability and validity of its findings. Here, we use DREAM to decompose the fast (*T*_*R*_ = 720ms) rfMRI data from the Human Connectome Project (HCP) [33] test-retest dataset, to 1) map the ranks of ALFF values through Slow-1, Slow-2, Slow-3, Slow-4, Slow-5 and Slow-6 bands and 2) evaluate the test-retest reliability of the ALFF metrics in these different frequency bands.

### Participants and Data Acquisition

The test-retest dataset from HCP consisting of 45 subjects were used for this analysis. All subjects were scanned with an HCP-customized Siemens 3T scanner at Washington University, using a standard 32-channel receive head coil. Three participants were excluded from the substantial analyses because their resting-state scan durations were shorter than others. Forty-two subjects (aged 30.3 ± 3.4 years, 29 males) were included in the present study. Each subject was scanned two times and each scan contained structural images (T1w and T2w), two rfMRI, seven runs of task fMRI and high angular resolution diffusion imaging (see details of the imaging protocols from HCP website). In the present work, we only used the rfMRI data, which consisted of 1200 volumes (*T*_*R*_ = 720 ms; TE = 33.1 ms; flip angle = 52°, 72 slices, matrix = 104 × 90; FOV = 208 × 180 mm; acquisition voxel size = 2 × 2 × 2 mm). The data were preprocessed according to the HCP MR preprocessing pipeline [13], resulting in the preprocessed surface time series data fed to the following DREAM analysis.

### Amplitude Analysis

For each rfMRI scan, we first extracted the representative time series for each of the 400 parcels [31] by averaging all the preprocessed time series within a single parcel. DREAM decomposed the time series into its components across the potential frequency bands. We performed ALFF analysis for all the bands of each run and each subject according to [47] implemented by CCS [36]. Subject-level parcel-wise ALFF maps for each frequency band were standardized into subject-level Z-score maps (i.e., by subtracting the mean parcel-wise ALFF of the entire cortical surface, and dividing by the standard deviation). The two standardized ALFF maps in the same session were then averaged, resulting in two (test versus retest) standardized ALFF maps per frequency band for each subject. To investigate the test-retest reliability of ALFF across the five frequency bands, we calculated the parcel-wise intraclass correlation (ICC) based upon the two ALFF maps [35,49]. We averaged the two standardized ALFF maps of all the subjects to obtain the group-level standardized ALFF maps. In order to evaluate the spatial distribution of the ALFF values for each parcel, we assigned its rank of ALFF values to the parcel (from 1 to 400). All the above analyses were done for each of the five frequency bands, leading to an ALFF ranking map for each frequency band.

### Results

DREAM decomposed the rfMRI timeseries into six frequency bands (Slow-6: 0.007 - 0.012 Hz; Slow-5: 0.012-0.030 Hz; Slow-4: 0.030-0.082 Hz; Slow-3: 0.082-0.223 Hz; Slow-2: 0.223-0.607 Hz; Slow-1: 0.607-0.694 Hz). Spatial rankings on ALFF are mapped in Figure 9. It is noticed that ALFF spatially ranked from high in ventral-temporal areas to low in ventral-occipital areas when the frequency band increased from low to high, while those in part of parietal and ventral frontal regions were reversed. The top-10 parcels are listed below:

**Figure 9.**
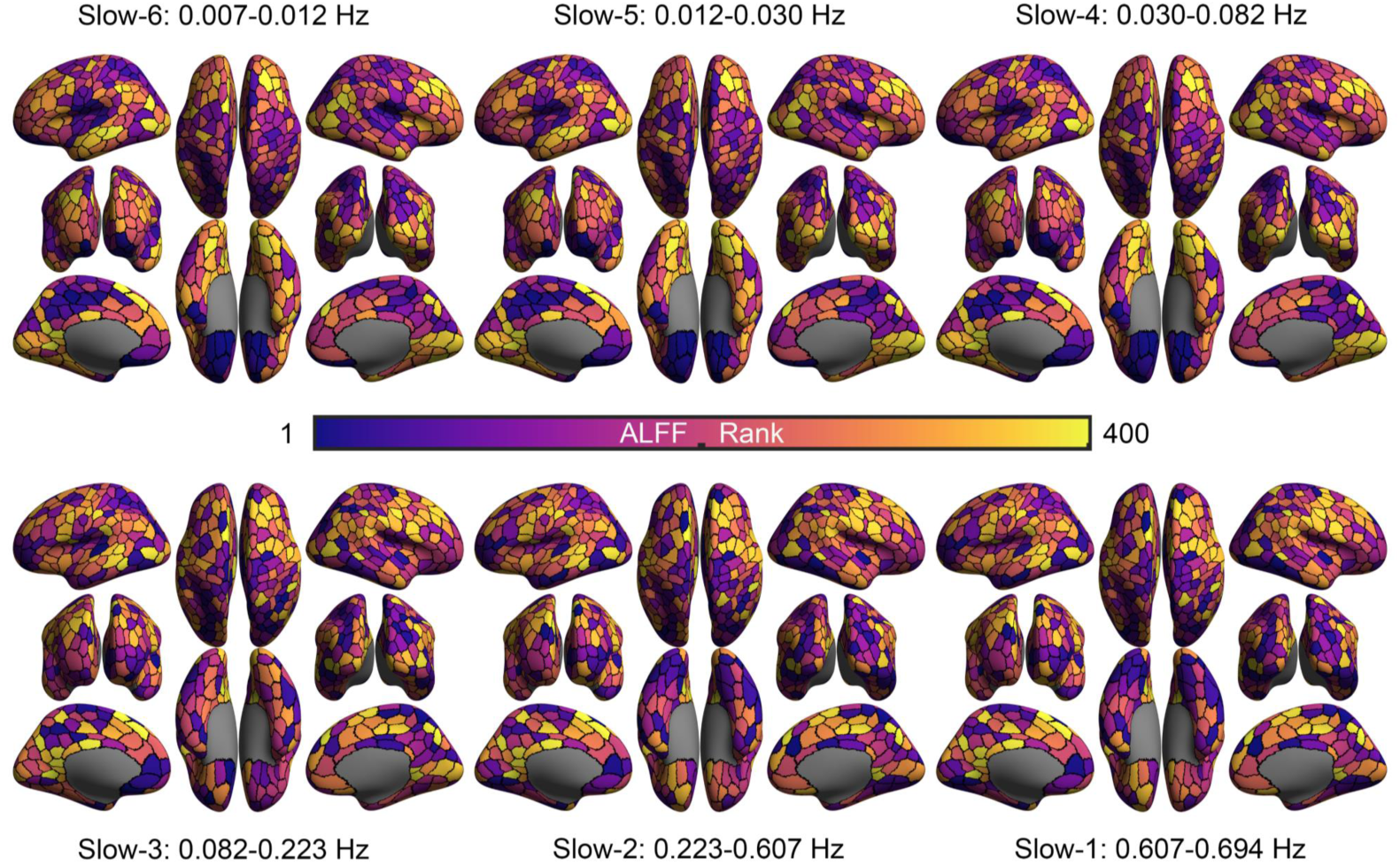
Spatially ranking ALFF across six frequency bands. **LH:**left hemi- sphere; **RH:** right hemisphere; Vis: visual network; SomMot: somatomotor network; DorsAttn: dorsal attention network; SalVentAttn: salience ventral attention network; Cont: frontal parietal control network; Default: default network; Limic: limbic network; see details of parcel naming at GitHub for the parcellation.

- Slow-6: LH_Default_pCunPCC_1, LH_Default_PFC_24,_RH_Default_PFCdPFCm_9, LH_Vis_16, RH_Vis 6, RH_Default PFCdPFCm_10,_LH Default Temp_7, LH_Default Temp_6, RH_Default_Par 3, _LH Default_pCunPCC 2
- Slow-5: LH_Default_PFC_24, RH_Default_PFCdPFCm_9, LH_Default_pCunPCC_1, LH_Vis_16, RH_Vis_6, LH_Vis_17, LH_Vis_5, RH_Vis 16, RH_Vis_15, RH_Default_PFCdPFCm 10
- Slow-4: LH_Vis_16, RH_Default_PFCdPFCm_9, LH_Default_pCunPCC_1, LH_Default_PFC_24, LH_Cont_PFCmp_1, RH_Cont_pCun_2, RH_Cont pCun_1, LH_Vis_17, _LH_Cont_Cing_2, RH_Vis_17
- Slow-3: LH_Vis_16, RH_Default_PFCdPFCm_9, LH_Default_PFC_24, RH_Cont_pCun_1, LH_Default_pCunPCC_1, LH_Cont_PFCmp_1,_RH_Cont Par_4, RH Vis_14, LH_Cont_Cing_2, _LH_Default_pCunPCC_5
- Slow-2: LH_Vis_16, RH_Default_PFCdPFCm_9, _RH_Cont_Par_4, LH_Default_PFC_24, RH_Cont_pCun_1, RH_Vis_14, LH_Cont_PFCmp_1, LH_Default_pCunPCC_1, LH_Default_pCunPCC 5, _LH_Cont_Cing_2
- Slow-1: LH_Vis_16, RH_Default_PFCdPFCm_9, RH_Cont_Par_4, RH_Cont_pCun 1, _LH_Vis 14, LH_Default_PFC 24,_LH_Cont_PFCmp_1, LH_Default_pCunPCC_1, LH_Default_pCunPCC_5, LH_Cont_Cing_2

Test-retest reliability maps of ALFF are also generated (Fig. 10) by mapping ICC using the linear mixed models. It is clear that the higher frequency bands, the more reliable ALFF measurements. The slow-2 (0.223-0.607 Hz) demonstrated the highest test-retest reliability of ALFF across the six frequency bands. The top-10 most reliable parcels are listed below:

**Figure 10.**
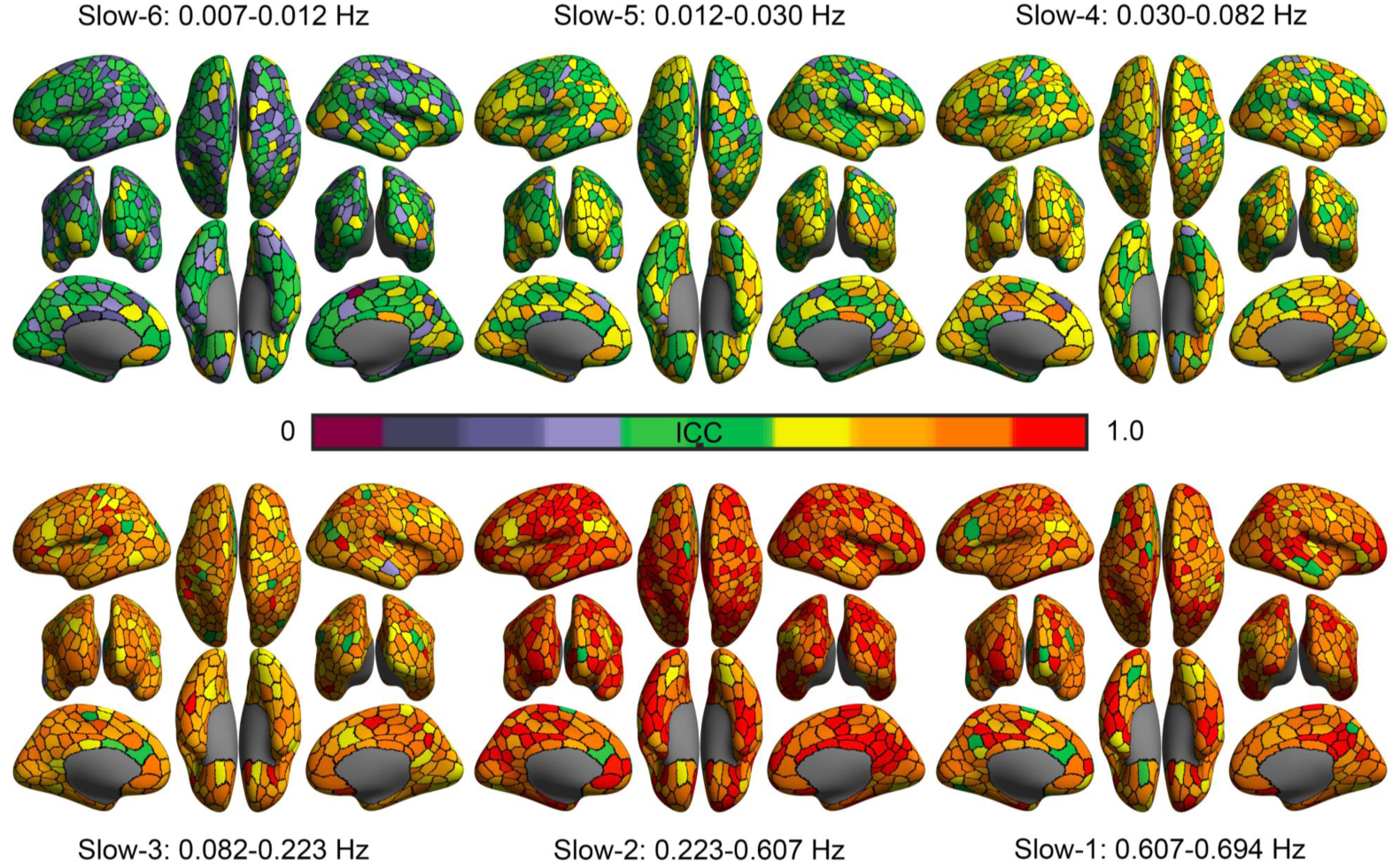
Test-retest reliability of ALFF across six frequency bands.

- Slow-6: RH_Vis_16, RH_Default_pCunPCC_5, RH_Cont_Cing_1, LH_Default_PFC_4, LH_Cont_PFCl_1, LH_Vis_14, RH_Vis_19, RH_Default_Temp_3, RH_Default_Temp_2, RH_Cont_PFCl_6
- Slow-5: RH_Cont_Cing_1, LH_Vis_23, LH_Default_pCunPCC_7, RH_Default_Temp_2, LH_Cont_OFC_1, RH_Cont_PFCl_15, RH_Default_pCunPCC_5, LH_Vis_14, RH_DorsAttn_Post_3, RH_Default_pCunPCC_9
- Slow-4: LH_Cont_OFC_1, RH_Default_pCunPCC_5, RH_Default_pCunPCC_9, RH_Cont_PFCl_15, LH_SalVentAttn_Med_1, RH_Cont_Par_5, LH_Cont_PFCl_2, RH_Cont_PFCv_1, RH_DorsAttn_Post_14, RH_Default_PFCv_1
- Slow-3: RH_SomMot_10, LH_DorsAttn_Post_1, RH_Default_PFCv_1, LH_Cont_Par_6, LH_SomMot_22, LH_Default_PFC_7, RH_DorsAttn_Post_14, RH_Default_pCunPCC_5, RH_DorsAttn_Post_3, LH_SalVentAttn_ParOper_1
- Slow-2: RH_Default_PFCv_1, LH_DorsAttn_Post_2, RH_Default_Temp_8, RH_Default_pCunPCC_5, LH_Default_pCunPCC_9, RH_Cont_PFCl_4, RH_Default_pCunPCC_6, RH_Default_pCunPCC_4, LH_Cont_Par_6, RH_Default_pCunPCC_7
- Slow-1: RH_Default_PFCv_1, RH_Default_pCunPCC_4, RH_Default_Temp_8, RH_Default_pCunPCC_8, RH_DorsAttn_Post_3, LH_Limbic_TempPole_4, LH_Cont_Par_6, RH_Default_pCunPCC_6, LH_DorsAttn_Post_1, LH_Default_PFC_6

## 5 Discussion

DREAM is a free and publicly available software that can decode oscillation data into multiple frequency bands. The simple interface was designed to allow all users to easily perform multi-band frequency analyses. The computational methods employed in DREAM to calculate the numbers and ranges of decoded frequency bands apply the Nyquist-Shannon sampling theorem and the brain oscillation theory [6]. Such a theory has been proven of great potentials to understand the brain dynamics as well as their behavioral correspondences. From a theoretical perspective, the oscillation theory can be independent of any modalities (e.g., EEG, MEG, ECoG, TMS, fMRI, fNIRS, eye tracking, etc.) for measuring these oscillations as windows into brain waves [3].DREAM is thus applicable for multiple forms of discrete sampling data, as long as the data are entered in the supported format. Currently, DREAM can process both NIFTI formatted neuroimaging data and text file formatted behavioral data while more other formats will be supported in its forthcoming releases.

As a demonstration of its utility, the results derived with DREAM for pure behavioral recordings suggest that head motion may be a behavioral feature reflecting both state and trait of individuals. We showed that head movements in the high frequency bands are more evident than those in the low frequency bands. This could be a behavioral reflection of the hierarchical organization of brain oscillations for their synchronization at multiple scales in space. Neural oscillations of the higher frequency-bands are related to more local information processing while the lower frequency-bands are for more distant communications in the brain. Our findings are consistent with the previous observation that the head motion had more impacts on the short-distance brain connectivity. While the head motion during fMRI scanning has been treated an important confounding factor in the neural signal [27], some recent work also argued its neurobiological components related to individual traits of the motor behaviors (e.g., [41,43]). The current researh offers data for an alternative explanation on such neurobehavioral trait likely driven by brain systems operating within a multi-band frequency landscape. In the context of development, as we expected, younger children moved more than older children across all the slow frequency bands. The stability of head motion during the experiment also varied with age, with head motion becoming less variable or more stable in older children. This is more evident in higher frequency bands, an implication that more sudden and sharp movements in younger children. Moreover, in a specific age range (7 - 9 years), boys moved more than girls across Slow-6 to Slow-1 bands but such differences vanished in the delta frequencies. This age range is a critical period for developing the ability to apply effective cognitive control (i.e., cognitive flexibility during executive function) [1], and our findings might reflect the sex differences in the cognitive development. In summary, our results demonstrate the necessity to study the frequency-specific characteristics of head motion, especially a perspective on understanding the neurobiological mechanism behind these behavior-related oscillations. This is of great potential to enrich our knowledge on the lifespan development such as children, the elderly and patients with neurologic or psychiatric conditions where both distance-related brain and the head motion measurements have been observed to correlate with each other [2, 10, 11, 30].

Differences in head motion across ages or between cohorts may reflect differences of certain traits, which may co-vary with detected brain signals and behavioral outcomes. The different properties of head motion in different frequency bands show that there may be different mechanisms associated with different frequencies. Head motion at higher frequencies varies more with age, and this may reflect that cognitive control associated with higher frequencies develops better with age. Of note, interpolation analyses indicated that this observation is not related to an issue of better signal-to-noise ratio at higher frequencies because there are more events per unit time. Within the narrow age range of 7 to 9 years old, boys moved more than girls in most frequency bands, although sex differences were larger at lower frequencies. This may indicate that the development of controlling system associated with lower frequencies may have larger sex-related differences for this age range. The above results lead us to speculate that there may be two control systems that are associated with different frequency bands of head motion which develop differently with age and between boys and girls. More detailed experimental studies deserved to test this postulation in future. The strategies of dealing with head motion issues in human brain mapping may also need updates regarding its measurement reliability and validity in terms of the possible neurobiological correlates [35,46,50]. One promising direction is to separate various sources of the head movements by using additional recordings or developing novel motion metrics (e.g., the recent progress in [24–26]). These efforts identified seven kinds of in-scanner motion in resting-state fMRI scans, and five of them related to respiration. Some pseudomotion occurred only in the phase encode direction and was a function of soft tissue mass, not lung volume. Using the Mock scanning experimental design as in the present work, together with the aforementioned approaches, could be of high value in further understanding neurobiobehavioral underpins of the human head movements.

Using fast fMRI from HCP, at the first time, we revealed the spatially configuring pattern of ALFF ranking gradually from low to high frequency bands. This indicates a trend along the two orthogonal axes. Along the dorsal-ventral axis, higher ALFF ranks were moving from the ventral occipital and the ventral temporal lobe up to regions in the parietal lobe as the frequency increasing. Along the anterior-posterior axis, from lower to higher bands, higher ALFF ranks were walking from the posterior to the anterior regions in the ventral part. This frequency-dependent ALFF pattern is similar to the findings of previous studies on the association between brain structure and gene expression, which also reported orthogonal gradations of brain organization and the associated genetic gradients [7,18]. The underlying physiological mechanism and functional significance of the frequency-dependent ALFF patterns deserve further investigations. It is interesting that the frequency-dependent pattern of ICC is quite uniform across the brain and as the frequency increased, its reliability increased alongside. This observation illustrated that compared with the low frequency bands, higher frequency bands might be more suitable for detecting individual differences in ALFF. Most of the previous studies have adopted ALFF of the lower frequency bands (i.e., Slow-5 and Slow-4 or around 0.01 to 0.1 Hz) where their ICCs rarely met the reliability requirement (*ICC* ≥ 0.8) of clinical applications. In contrast, our findings suggest that both Slow-2 and Slow-1 ALFF could be the usable and reliable marker of the brain oscillations for these applications. It is noticed that the reliability of Slow-1 ALFF is slightly lower than those of Slow-2 ALFF, and this may be an indication on the limited Slow-1 band here compared to its theoretical range (around 0.6065 − 1.6487 Hz). While studies of the very fast sampled fMRI signals such as HCP are sparse, it is quite promising for future studies with multiple neuroimaging modalities (e.g., [3,15]) to DREAM as an integrative tool across frequencies. An open toolbox such as DREAM is essential for large-scale projects inspired by the increasing practice of open sciences coming with more and more fMRI and EEG datasets openly shared as well as their applications (e.g., [45]).

## Information Sharing Statement

The DREAM toolbox is fully open to the public by sharing both the off-line version (https://github.com/zuoxinian/CCS/tree/master/H3/DREAM) and the light online version (http://ibraindata.com/tools/dream). To ensure the reproducibility of our findings, all the codes and head motion data for generating the figures and other results in the present work are also shared via DREAM and CCS website.

- Connectome Computation System: https://github.com/zuoxinian/CCS
- DREAM: https://github.com/zuoxinian/CCS/tree/master/H3/DREAM
- Visualization Data in DREAM1 (GraphPad): github.com/zuoxinian/CCS/blob/master/H3/DREAM/DREAM1_demo.pzfx
- ANOVA Codes in DREAM1 (MATLAB): github.com/zuoxinian/CCS/blob/master/H3/DREAM/DREAM1_repANOVA.m

Please credit both DREAM and CCS work if you use our DREAM in your research.

## Acknowledgments

This work was supported in part by the China - Netherlands CAS-NWO Programme (153111KYSB20160020), Beijing Municipal Science and Technology Commission (Z161100002616023, Z171100000117012), the National R&D Infrastructure and Facility Development Program of China, Fundamental Science Data Sharing Platform (DKA2019-12-02-21), and Guangxi BaGui Scholarship (201621). The neuroimaging data were provided by the HCP WU-Minn Consortium, which is funded by the 16 NIH institutes and centers that support the NIH Blueprint for Neuroscience Research 1U54MH091657 (PIs: David Van Essen and Kamil Ugurbil), the McDonnell Center for Systems Neuroscience at Washington University.

## Contribution

Conceived and designed the experiments: Xi-Nian Zuo, Hai-Fang Li. Performed the experiments: Zhu-Qing Gong, Peng Gao, Hao-Ming Dong, Chao Jiang, Xi-Nian Zuo. Analyzed the data: Zhu-Qing Gong, Peng Gao, Chao Jiang, Xi-Nian Zuo. Contributed reagents/materials/analysis tools: Zhu-Qing Gong, Peng Gao, Chao Jiang, Xi-Nian Zuo. Wrote the paper: Zhu-Qing Gong and Xi-Nian Zuo drafted the manuscript and all the authors revised the manuscript.

